# Differential and regionally distinct molecular responses distinguish FLASH and conventional RT in a medulloblastoma model while preserving therapeutic outcomes

**DOI:** 10.64898/2026.03.16.712103

**Authors:** Michèle Knol, Javier Franco-Pérez, Aymeric Almeida, Louis V. Kunz, Benoit Petit, Antoine Job, Jonathan Ollivier, Jackeline Romero, Jeannette Jansen, Veljko Grilj, Charles L. Limoli, Marie-Catherine Vozenin, Paola Ballesteros-Zebadúa

## Abstract

**Background:** FLASH radiotherapy might minimize treatment-related complications in medulloblastoma patients. To support its clinical translation, we investigated the therapeutic index of hypofractionated FLASH- and CONV-RT and their respective molecular determinants using an orthotopic human medulloblastoma mouse model.

**Methods:** Five cohorts of juvenile UW228-MB-bearing nude mice were irradiated or not using 3×10 Gy (BED_10 =_ 60 Gy), delivered 48 h apart at 0.1 Gy/s (CONV) or 5.5×10⁶ Gy/s (FLASH) using an electron beam (eRT6). Digital spatial profiling (DSP) and immunofluorescence were performed 24h after radiotherapy in one cohort, while four cohorts were followed for long-term tumor response, cognition and neuroinflammation.

**Results:** Both CONV- and FLASH-RT induced a long-lasting anti-tumor response in all irradiated animals. Long-term NOR performance was preserved in irradiated animals, with no significant difference compared with age-matched, non-tumor-bearing controls. A higher but non-significant proportion of very good discriminators was observed in the FLASH cohort while neuroinflammatory markers were statistically reduced. In the FLASH-irradiated mice, DSP revealed a sustained microglial activation in the cerebellar tumor micro-environment, with enhanced expression of genes with phagocytic and proteolytic activity. In the hippocampus, FLASH-induced genes involved a preferential astrocyte/neuron transcriptional crosstalk including metabolic support and synapse-glia interaction. The relevance of this metabolic support provided by astrocytes to neurons in the FLASH cohort was further supported by the enhanced expression of the astrocytic monocarboxylate transporter MCT4, a key transporter of lactate.

**Conclusion:** The study shows that despite similar therapeutic index, hypofractionated FLASH and CONV-RT induce a distinct molecular signature in a human medulloblastoma mouse model.

**Key Points:**

- Similar therapeutic outcome after hypo-fractionated FLASH and CONV.
- FLASH reduced neuroinflammation
- FLASH induced clearance-prone phagocytic and proteolytic activity in the microglia of the TME.
- FLASH programs a neuro-supportive astrocyte phenotype in the hippocampus.

**Importance of the study:** This study constitutes a milestone for the implementation of FLASH-RT for the treatment of children with brain cancer, demonstrating that hypo-fractionated (3x10 Gy) FLASH- and CONV-RT are equally tumor-ablative and sufficient to preserve long-term NOR performance. Despite the equivalence of therapeutic outcomes, FLASH- and CONV-RT trigger fundamentally distinct biological responses. FLASH promotes a metabolically active, phagocytosis-prone phenotype in microglial cells consistent with immune activation and tumor surveillance. In contrast, CONV promotes proliferative and immunosuppressive signaling programs. Interestingly, FLASH also differentially shapes long-term transcriptional programs in hippocampal subregions critical for memory encoding, pattern separation, and consolidation, while reprogramming astrocytes toward a neuro-supportive phenotype. Collectively, these findings reveal that equivalent tumor control does not imply biological equivalence: FLASH engages a distinct brain response characterized by coordinated immune surveillance and neuro-supportive programs. Data supports the potential of translating FLASH-RT to cure pediatric brain tumors while preserving the functional integrity of the developing brain.

## Introduction

The treatment of medulloblastoma (MB) is based upon biological and pathological MB subtypes, molecular biomarkers, staging, age at diagnosis and possible genetic predisposition. Multimodal treatment is composed of initial maximal safe surgical resection followed by craniospinal irradiation (CSI) of 23.4 Gy in 13 fractions, followed by a boost to the tumor bed (30.6 Gy in 17 fractions) for a total dose of 54 Gy, following the PNET 5 protocol (NCT02066220) for standard-risk medulloblastoma. Meanwhile, high-risk medulloblastoma patients receive CSI of 36 Gy in 20 fractions, with a boost (18 Gy in 10 fractions) for a total dose of 54 Gy and intensive chemotherapy regimens^1^. Sites of metastasis may receive additional RT boosts. While tumor control is excellent, with 80% of patients being cured, one major limitation of radiotherapy in children is the extreme vulnerability of the developing brain and the relative radiation resistance of brain tumors. The immature central nervous system is highly radio-sensitive, and the standard fractionation regimen used, composed of 1.8–2.0 Gy per fraction, is known to impair neurogenesis, synaptic development, and white matter maturation, ultimately leading to long-term neurocognitive deficits, including reduced IǪ, memory impairment, and executive dysfunction^2, 3^. The survivors and their families therefore face challenges related to long-term side effects affecting their quality-of-life. The brain of young children is also known to have a lower tolerance to larger fraction sizes, limiting the possibility to apply altered hypofractionation regimen possibly more efficient against tumors. A last practical aspect is repeated anesthesia needed for the immobilization of the children during daily fractionated radiotherapy, introducing operational complexity both for health professionals and families, as well as medical risks associated with repeated sedation.

A recent meta-analysis showed that CSI using protons provides equivalent survival to photon-based regimen while significantly reducing growth hormone deficiency (GHD), hypothyroidism, mild ototoxicity, and neurocognitive toxicities in children with medulloblastoma^4^. However, novel therapeutic strategies able to further spare healthy brain and reduce treatment burden are needed. A decade ago, we pioneered FLASH radiotherapy^5, 6^, which can provide a unique opportunity to enhance therapeutic outcome by minimizing complications. In experimental adult and juvenile animal models, most studies using electron, photon and proton FLASH have demonstrated reduced cognitive decrements while maintaining anti-tumor efficacy^7–13^, even when hypo-fractionated regimen were used^14, 15^, except one^16^. Recently, the combination of a single 10 Gy dose of proton-FLASH with GD2-CAR-T cell therapy was shown to enhance survival of MB transgenic SmoM2 mice^17^.

The current study was designed to provide key missing elements to support the clinical translation of FLASH for the treatment of children with brain tumors. A clinically relevant hypo-fractionated radiotherapy regimen (biologically effective dose (BED_10_ of 60 Gy) achieved robust tumor ablation in an orthotopic human medulloblastoma model while mitigating radiation-associated brain injury in a dose-rate independent manner. Despite these equivalent therapeutic outcomes, molecular responses were dose rate dependent. Cerebellar transcriptomic analyses revealed that microglial cells are the principal acute responders to irradiation, with FLASH promoting a metabolically active, phagocytosis-prone phenotype, while proliferative and immunosuppressive signaling programs were induced by CONV. In the adjacent healthy hippocampus, FLASH induced a coordinated astrocyte-to-neuron transcriptional program rather than a uniform inflammatory response. While our radiotherapy regimens were not observed to induce significant cognitive decline, at the molecular level FLASH induced astrocytic reprogramming and reduced neuroinflammatory footprint able to support neuronal function in three subregions known to be critical for memory encoding, pattern separation, and consolidation.

## Material and Methods

### Human medulloblastoma model in mice

Swiss nude mice (NU(Ico)-Foxn1nu) were bred in house and both juvenile female and male mice were included in five successive cohorts leading to variable number of animals and sex representation in each cohort (Fig. 1A; Supp Fig. 2A). Tumor growth, survival and behavior were evaluated in both sexes, the non-tumor-bearing control group used for behavior, spatial transcriptomic, immunofluorescence and long-term neuroinflammatory analyses were performed in females only as no males were available. Experiment was approved by the Swiss (VD3603) Ethics Committee for Animal Experimentation and performed in accordance with institutional guidelines.

**Figure 1.**
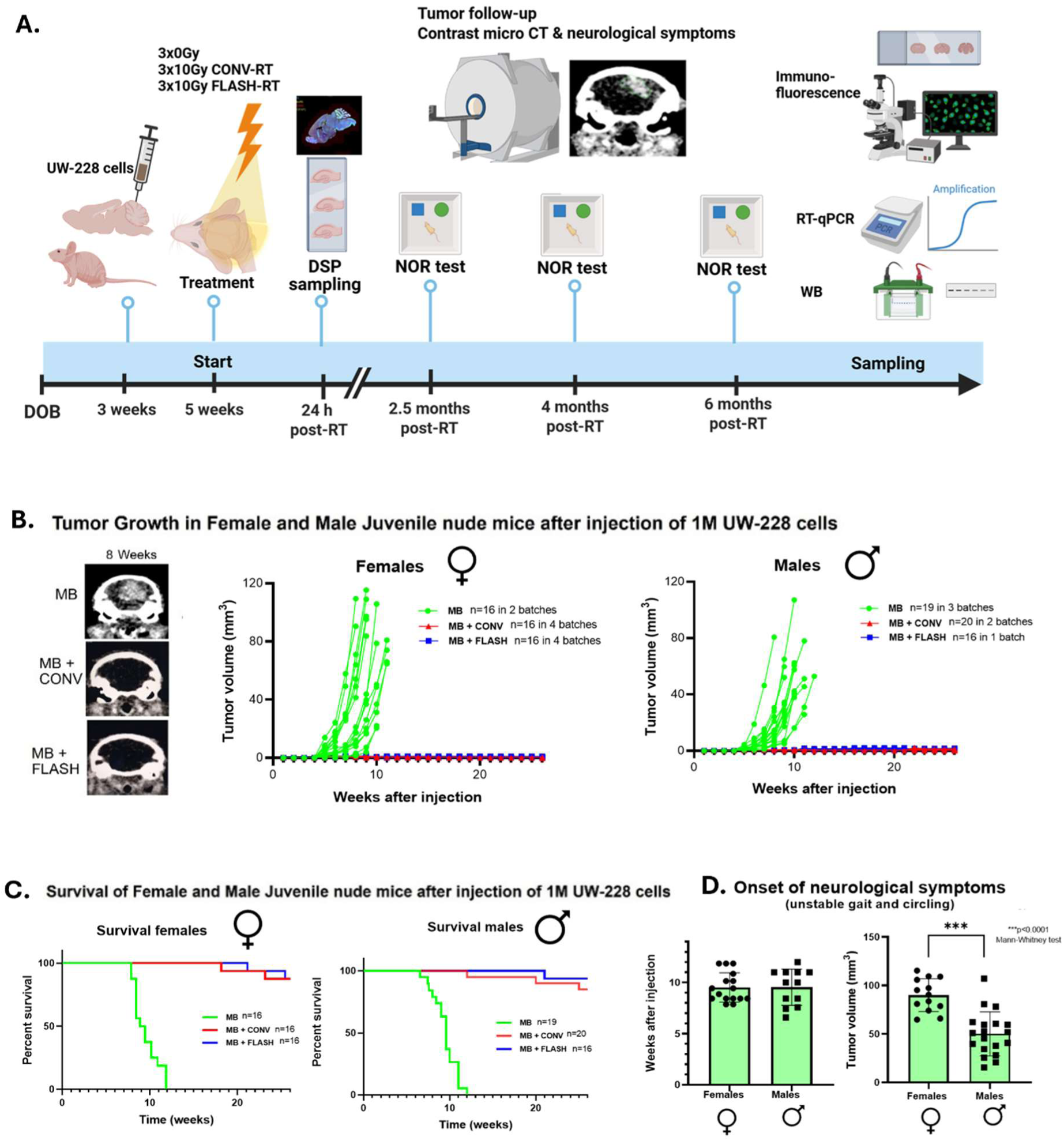
Long-term anti-MB response following FLASH- and CONV-RT in medulloblastoma-bearing mice. **A-**Diagram showing the timeline of the experiments. All animals were in good health at the time of behavioral studies, and sampling was done in accordance with ethical procedures. **B-**Tumor volume was evaluated weekly for 26 weeks for both male and female cohorts and example images obtained by micro-CT showing tumor growth 8 weeks after MB cell implantation showing an isoefficient control of MB after FLASH-RT and CONV-RT. **C-**Survival curves of MB-bearing mice after whole brain irradiation with a fractionation schedule of 3x10 Gy with FLASH-RT or CONV-RT for both sexes. The Kaplan–Meier curves represent overall survival and include events not demonstrated to result from tumor progression. **D-**Control MB mice exhibited neurological symptoms at a median of 9-10 weeks after injection for both sexes. Median tumor volume at the onset of neurological symptoms was 90.4 mm³ for females and 51 mm³ for males and was statistically different (** p<0.0001 Mann-Whitney test).

**Figure 2.**
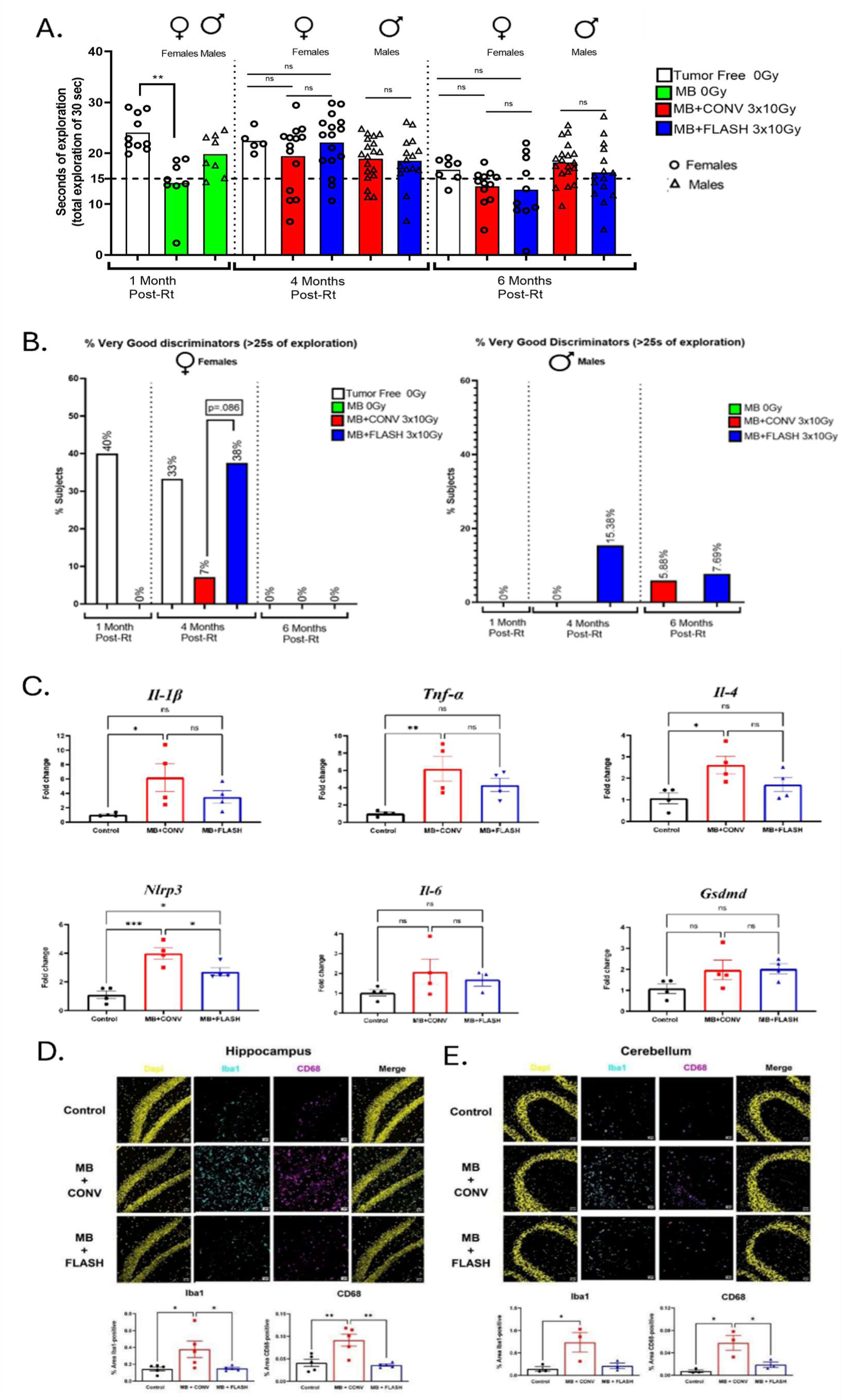
Long-term novel object recognition and neuroinflammatory response following FLASH- and CONV-RT in medulloblastoma-bearing mice. **A–**Novel object recognition (NOR) quantified as the time spent exploring the novel object during the first 30 s of cumulative object exploration at 1, 4, and 6 months post-radiotherapy (RT). The dashed lines indicate 15 s (50% of total exploration time) and 25 s, the threshold used to classify mice as very good discriminators. Female cohorts included tumor-free non-irradiated controls, medulloblastoma-bearing non-irradiated mice (MB 0 Gy), and medulloblastoma-bearing mice treated with CONV-RT or FLASH-RT (3 × 10 Gy). Male behavioral data were available for MB 0 Gy, CONV-RT, and FLASH-RT groups; corresponding tumor-free male controls were not available. Independent experimental batches and their sex composition are detailed in Supplementary Figure X. Potential outliers were identified using the ROUT method (Ǫ = 5%). Continuous behavioral outcomes were analyzed using the Kruskal–Wallis test followed by Dunn’s multiple-comparison test. Bars represent mean ± SEM and individual values are shown; circles indicate females and triangles indicate males. B–Exploratory categorical analysis showing the percentage of female (left) and male (right) mice classified as very good discriminators, defined as spending >25 s of the first 30 s of cumulative object exploration exploring the novel object. CONV-RT and FLASH-RT proportions were compared using two-sided Fisher’s exact tests. In females, the difference at 4 months post-RT did not reach statistical significance (*p* = 0.086); corresponding comparisons in males were also non-significant (4 months, *p* = 0.1573; 6 months, *p* > 0.9999). C–Gene expression of the inflammatory markers *Il-1b, Tnf-a, Il-C, Nlrp3, Il-4,* and *Gsdmd* in whole-brain tissue from control mice and mice treated with CONV-RT or FLASH-RT. Data are expressed as mean ± SEM (*n* = 4 per group) and were analyzed by one-way ANOVA followed by Tukey’s post hoc test. *p* < 0.05, **p < 0.01, ***p < 0.001. D, E–Immunofluorescence staining of Iba1 and CD68 in the hippocampus (D) and cerebellum (E), showing increased microglial activation following CONV-RT but not FLASH-RT.

The UW228 cells were obtained from a 9-year-old female patient thanks to a donation from Prof. Martin Baumgartner, University Children’s Hospital Zürich, authorized by Dr. John Silber C Michael Bobola from Seattle Children’s Hospital and Regional Medical Center. The α/β ratio of the UW228 cell line, estimated from clonogenic survival data using the linear-quadratic model, was 22.1 Gy (Supp. Fig. 1A). UW228 human MB cells were cultured in a complete medium containing DMEM + GlutaMAX (31966–021, Gibco, USA) and supplemented with 10 % FBS (F7524, Sigma-Aldrich, USA). 1 million UW228 cells were engrafted in the cerebellum of juvenile mice right after the weaning (3 weeks of age), according to the following stereotactic coordinates: AP -6, ML +1, DV +2, at an angle of 30 degrees.

### Hypo-fractionated radiotherapy

Animals were divided into four experimental groups: sham-irradiated tumor-free controls, sham-irradiated tumor-bearing controls, and tumor-bearing mice treated with conventional radiotherapy (CONV), or FLASH radiotherapy. The radiotherapy regimen was composed of three fractions of 10 Gy delivered 48 h apart under anesthesia.

Mice were irradiated two weeks after tumor engraftment at 5 weeks of age. Whole-brain irradiation was delivered with 6 MeV Oriatron eRT6 electron beam linear accelerator (PMB, Alcen, France) using a graphite 17 mm half circle collimator, protecting the eyes but covering the entire brain, including the cerebellum, as previously described^18^, and using daily dosimetric control^18^. Each fraction of 10 Gy was delivered with CONV at 0.1 Gy/s or with FLASH in a single pulse at 5.5x10^6^ Gy/s. All beam parameters are listed in Supp. Table 1.

### Tumor response

Tumor growth was monitored once per week using a micro-computed tomography (micro-CT) system at 55 kV, 0.19 mA, and 75 ms (MILabs, The Netherlands), using contrast enhancement (Accupaque 350 mg/mL; GE Healthcare AG, Switzerland). Tumor volumes were measured from DICOM files, performing slide-by-slide segmentation and an automated ellipsoidal volume formula to calculate the final volume. Additionally, HE staining was performed to confirm the presence of the tumor in MB-bearing mice 24h after radiotherapy (Supp. Fig. 1B).

### Novel object recognition

The novel object recognition (NOR) test in mice is widely used to investigate radiation-induced cognitive decrements^19^ and has been performed as described previously in FLASH studies^8, 9, 20^ however we modified the analytical part for several reasons: First, the T-cell deficiency of nude mice has been shown to modify hippocampal memory-related signaling ^21^. Second, tumor implanted Swiss nude mice frequently exhibited prolonged freezing and variable latency before initiating object exploration, therefore we analyzed their performance based on a fixed cumulative object-exploration criterion. Third, four cohorts of animals were analyzed sequentially (supp. Fig. 2A,B). For this analysis, the time spent exploring the novel object during the first 30 s of cumulative exploration of either object was quantified, irrespective of the total time required to reach 30 s of object exploration. This approach reduced the influence of inter-animal differences in freezing, locomotor activity, and exploration latency. Because cumulative object exploration was fixed at 30 s, novel-object exploration time is presented (Figure 2A). Age-matched non-tumor-bearing animals were included as controls when available. Statistical analyses were performed using GraphPad Prism (GraphPad Software, San Diego, CA, USA). Potential outliers were identified using the ROUT method (Ǫ = 5%) before assessment of normality. Normality was evaluated using the Shapiro–Wilk test. As normality was not satisfied across all behavioral groups, continuous behavioral outcomes were analyzed using the Kruskal–Wallis test followed by Dunn’s multiple-comparison test, with comparisons restricted to groups evaluated at the same time point. Male behavioral data were analyzed separately as the corresponding non-tumor-bearing male control group was not available. Experimental cohort composition and batch-specific behavioral data are provided in Supp. Fig. 2. In addition, mice spending >25 s of the first 30 s of cumulative object exploration exploring the novel object (>83.3%) were classified as very good discriminators. The proportions of very good discriminators in the various groups were compared using two-sided Fisher’s exact tests.

### GeoMx Digital Spatial Profiling

To gain insight into the transcriptional response at an acute timepoint after the different irradiation modalities, spatial transcriptomic was performed according to the standard nanoString GeoMx Digital Spatial Profiler (DSP) protocol^1^, using the Whole Transcriptome Atlas (WTA) panel on chosen brain areas. Brains from 3 mice per group (tumor-bearing control, CONV, and FLASH) were sampled 24 hours post-irradiation (Figure 1A). Regions-of-interest (ROI) were manually delineated in the cerebellum and in the dentate gyrus (DG), cornu ammonis 1 (CA1) and cornu ammonis 3 (CA3) of the hippocampus (Figure 3A). Two ROIs were delineated in the cerebellum.

**Figure 3.**
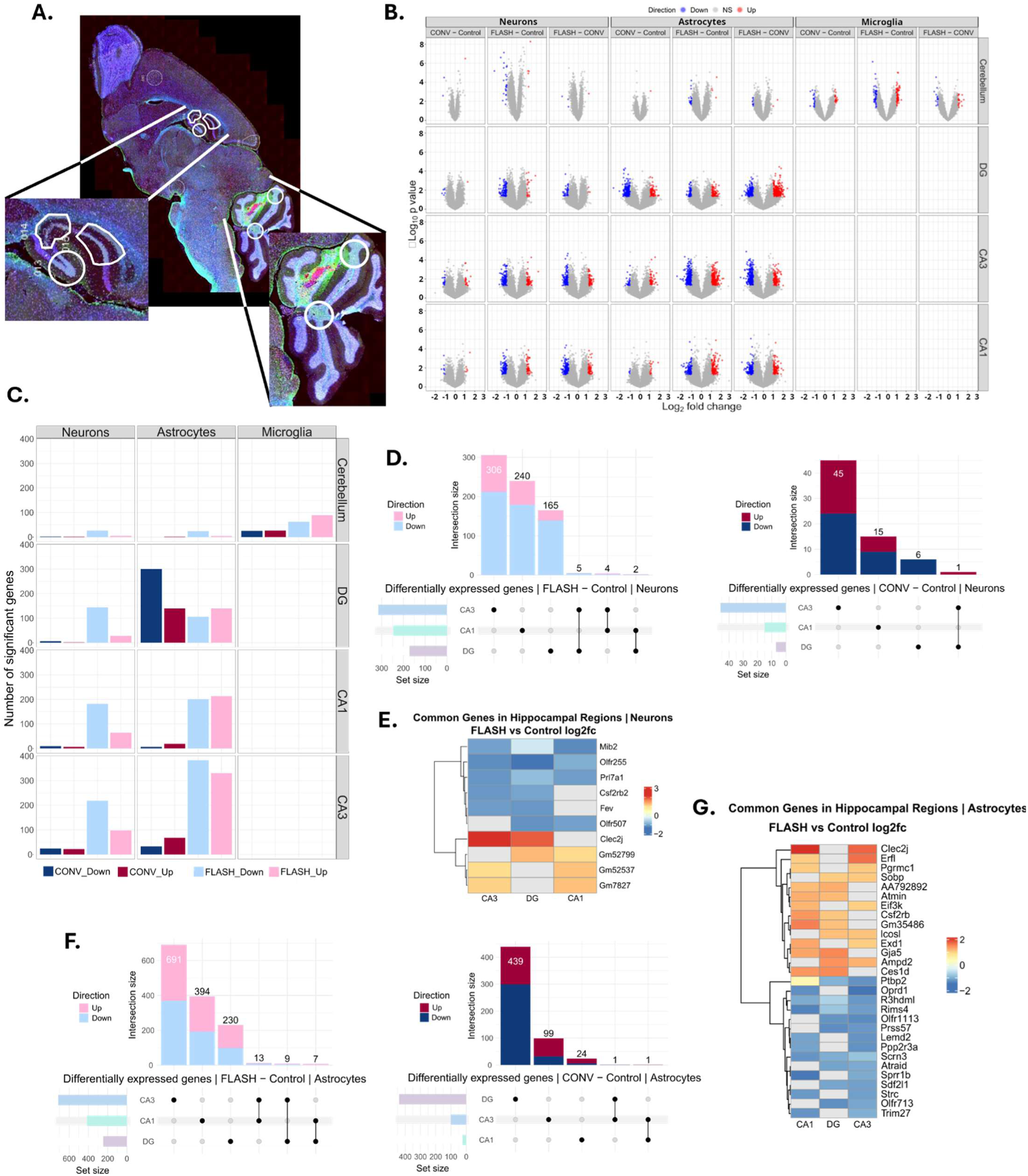
FLASH-induced transcriptomic response is region- and cell-specific. **A-**Representative image of a stained slide used for the DSP with the regions-of-interest in the cerebellum and the hippocampus (DG, CA1, and CA3). A 20x magnification was used. Nuclei are shown in blue, neurons in yellow, microglia in red, and astrocytes in green. **B-**Volcano plots showing the differential gene expression (DGE) analysis of all the regions and cell types. Cutoff values of |log2(fc)|>1 and p-value < 0.05 were used. **C-**Barplot of all significantly differentially expressed (DE) genes from the DGE, separated by region, cell type and irradiation modality. **D-**UpSet plots showing the intersections between the genes found significantly expressed in the different hippocampal regions in the neurons after FLASH (light blue and pink) and CONV (blue and red). **E-**A heatmap of all the common genes as found in D in the hippocampal regions for the neurons that were found to be differentially expressed after FLASH compared to control. Grey means that this gene was not significantly differentially expressed in that region. **F**-UpSet plots showing the intersections between the genes found significantly expressed in the different hippocampal regions in the astrocytes after FLASH (light blue and pink) and CONV (blue and red). **G**-A heatmap of all the common genes as found in D in the hippocampal regions for the astrocytes that were found to be differentially expressed after FLASH compared to control. Grey reflects no significantly differentially expressed genes in that region.

### Bioinformatics

Ǫuality control, pre-processing, and further analysis of the data was performed in R (v4.4.0) as described in the supplementary methods.

Differential gene expression (DGE) analysis was performed per region and cell type using a linear mixed model with the irradiation condition as a fixed effect and the slide as a random intercept to account for technical variability. Genes with an absolute log2 fold change >1 and a p-value of <0.05 were considered significantly differentially expressed. Gene set enrichment analysis (GSEA) was performed per region and cell type on gene lists ranked by the metric−(log2(fc)) ⋅ (−log10(*p value*)), combining the size of the effect and the significance. For further visualization, normalized expression values were used.

All scripts used for the analysis and plotting can be found at https://gitlab.unige.ch/lirr/dsp-pediatric-mb, as well as the raw sequencing data, which is available at https://www.ncbi.nlm.nih.gov/geo/ under accession number GSE325496.

### Immunofluorescence and q-RT-PCR

DSP samples (n=3 per group) were used to assess GFAP and MCT4 expression on serial sections 24h post-RT. Six months post-RT (n=6 per group) mice were sampled by intracardiac perfusion with PBS, and the brain was processed to analyze neuroinflammation. Specific procedures are described in the supplementary material and method.

## Results

### Both FLASH- and CONV-RT induce a durable anti-MB response

Juvenile mice underwent radiotherapy 2 weeks post-UW228 tumor engraftment. Radiotherapy induced complete and long-lasting response in 32/32 female and 36/36 male mice (Fig. 1B) transpiring in long-term overall survival (Fig. 1C) while 16/16 non-irradiated females and 19/19 males from 5 distinct cohorts developed MB visible on micro-CT imaging 8 weeks post-engraftment and developed neurological symptoms including unstable gating and circling within 9-12 weeks that required euthanasia (Fig. 1D).

### Long-term novel object recognition after FLASH and CONV-RT

Tumor-bearing, non-irradiated mice exhibited altered NOR performance before the development of neurological symptoms, showing that medulloblastoma growth impaired cognition (Fig. 2A). For long-term term follow-up analysis age-matched tumor-free female controls were included. NOR performance was similar in tumor-free controls, CONV- and FLASH-RT cohorts of females at 4 and 6 months post-RT (Fig. 2A), results that were likely associated with an age-related drop in performance across the groups. For males no tumor-free animals were available but again similar NOR performance was measured after CONV and FLASH-RT. The composition of each experimental cohort and batch-specific behavioral results is provided in Supp Fig. 2.

Next, we assessed each animal individually and found a higher proportion of very good discriminators in FLASH-irradiated animals than in CONV-RT in both sexes (Fig. 2B). At 4 months post-RT, 38% of FLASH-treated females were classified as very good discriminators and only 7% after CONV-RT (*p* = 0.086); 15.38% of FLASH-irradiated males vs 0% of CONV-irradiated were very good discriminators (*p* = 0.1573). However, statistical significance was not reached.

### Neuroinflammation is reduced after FLASH

At endpoint (6 months post-RT), age-matched tumor-free non-irradiated animals and irradiated animals were sampled to investigate neuroinflammatory status and glial activation in the cerebellum and the hippocampus. A panel of pro-inflammatory cytokines was analyzed by qRT-PCR on whole brain slices. *Il-1β, TNF-α*, IL4 and *Nlrp3* mRNA levels were found to be enhanced uniquely after CONV irradiation vs control, while the level of IL-6 and Gasdermin-d mRNA was not different in control and irradiated animals (Figure 2C). Both in the cerebellum and the hippocampus, high expression levels of *Iba1* and *CDC8* were measured in CONV-treated animals, suggesting glial activation whereas it was similar in FLASH-treated and tumor-free non-treated animals (Figure 2D, E). GFAP staining revealed no astrogliosis induction in the irradiated as compared to non-irradiated animals (Supp. Figure 4A, B).

**Figure 4.**
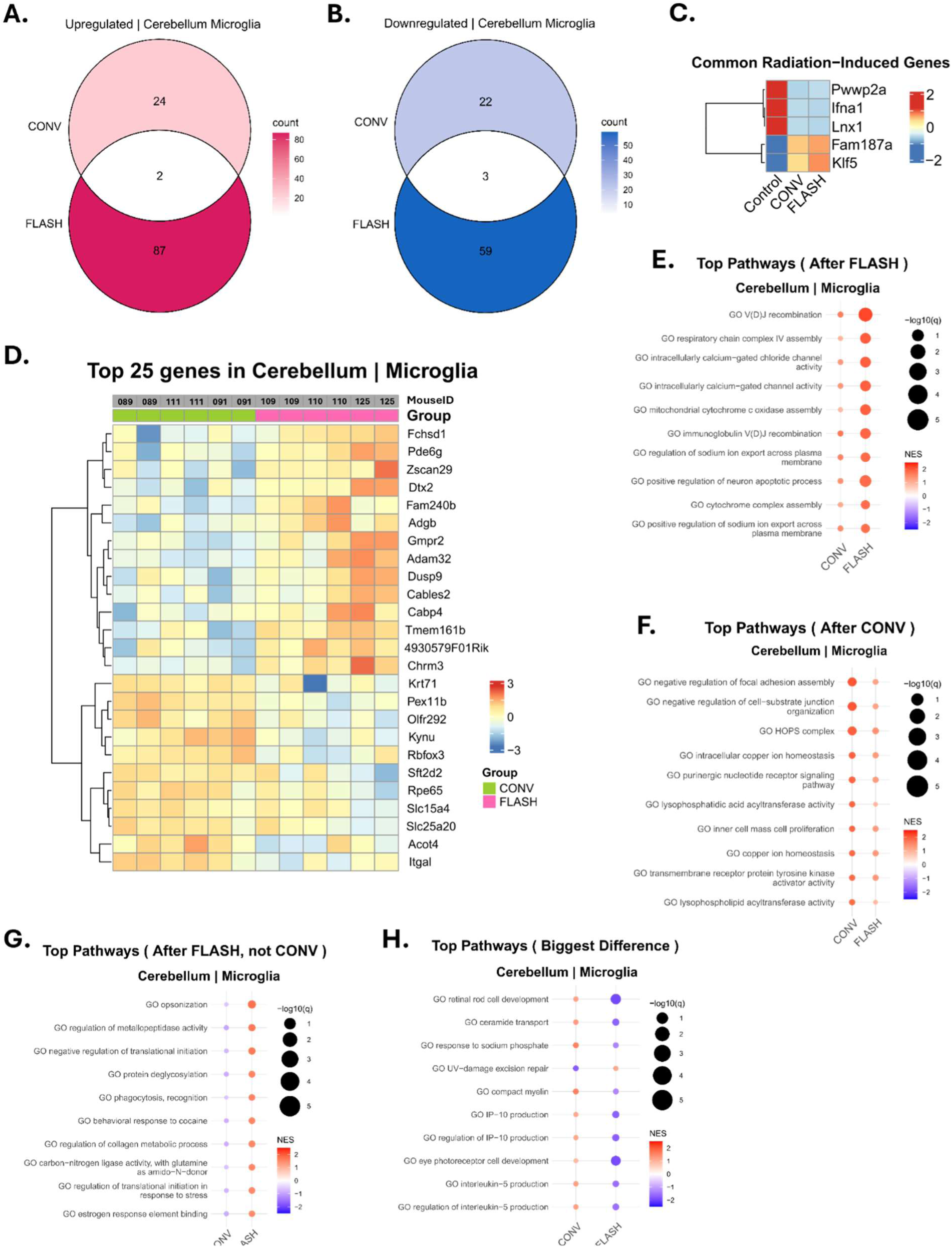
FLASH and CONV induce distinct microglial transcriptomic imprints in the cerebellum. **A-**Venn diagram of the intersection between FLASH and CONV of the upregulated significant DE genes in the microglia in the cerebellum. **B-**Venn diagram of the intersection between FLASH and CONV of the downregulated significant DE genes in the microglia in the cerebellum **C-**Clustered heatmap of the common radiation-induced DE genes, from the intersections of the Venn diagrams, averaged per group and scaled per row. **D-**Heatmap with the top 25 DE genes with the biggest difference between CONV and FLASH. Genes were ranked according to |log2fc|*-log10(p-value). Columns represent a segment, two ROIs in the cerebellum were used per mouse. Rows were hierarchically clustered. **E-**Top 10 pathways from the GSEA ranked by the highest absolute normalized enrichment score (NES) after FLASH, where a positive NES (red) represents an upregulated pathway and a negative NES (blue) a downregulated pathway. The size of the dots shows the log10(q-value), i.e. the false discovery rate. **F-**Like E, pathways were ranked by the absolute NES after CONV. **G-**Similar to E, pathways were ranked according to the highest absolute NES after FLASH when NES after CONV was negative. **H-**Similar to E, pathways were ranked according to the biggest difference between the absolute NES after CONV and FLASH.

### FLASH-induced transcriptomic response is region- and cell-specific

Differential gene expression analysis in various regions of the brain (Fig. 3A) showed more transcriptional changes after FLASH than after CONV (Fig. 3B). Interestingly, in the cerebellum, only microglial transcriptomic was modulated by radiotherapy and different between FLASH and CONV. Meanwhile, in the hippocampal regions, a radiation induced transcriptional profile was observed in astrocytes and neurons. We also found that FLASH induced specific transcriptional changes both in neurons and astrocytes in the DG, CA1 and CA3 subregions. These transcriptional changes in the neurons seemed to involve mostly downregulated genes, while in the astrocytes an equal proportion of down- and upregulated genes were found (Fig 3C). Contrary to that, CONV mainly altered the astrocytic transcription in the DG, while it had almost no transcriptional impact on the other regions or cell types at this timepoint (Fig. 3C). Furthermore, we observed that significantly differentially expressed genes were region-specific, while only selected few genes were common to multiple regions (Fig. 3D-E). Amongst those common genes was *Clec2j*, (C-type lectin like receptor 2), found to be upregulated after FLASH in the neurons of the CA3 and DG and the astrocytes of the CA1 and CA3 (Fig. 3E, G). Interestingly, *Clec2j* was also found to be upregulated in the neurons and astrocytes in the CA1 and CA3 after CONV. In the astrocytes of the CA1 and CA3 *Trh*, (tryrotropin-releasing hormone), was uniquely downregulated after CONV. Finally, *Mib2*, encoding a ubiquitin protein ligase, was downregulated after FLASH in the neurons of the CA1, CA3, and the DG (Fig. 3E).

### FLASH and CONV induce a distinct microglial transcriptomic signature in the TME

An early microglial signature was induced by irradiation in the cerebellum, while neurons and astrocytes were mostly silent (Fig 3B, C). A distinct profile was revealed after FLASH and CONV irradiation, where only 2 genes were commonly upregulated (*Fam187a* and *Klf5*) and 3 were downregulated (*Pwwp2a*, *Ifna1*, and *Lnx1*) (Fig. 4A, B, C). A total of 87 genes were found to be upregulated solely by FLASH, and 59 downregulated, meanwhile only 24 genes were uniquely upregulated by CONV and 22 downregulated (Fig. 4A, B, Supp. Table 3). The heatmap of the top 25 differentially expressed genes with the biggest difference between CONV and FLASH showed some variations in intensity within the different ROIs but nonetheless revealed a common signature in the microglial cells, containing many newly identified, radiation induced genes (Fig. 4D). Additional GSEA analysis revealed that the FLASH-induced pathways are related to VDJ recombination, calcium/chloride gated channels and sodium ion export, as well as mitochondrial function (Fig 4E), suggesting a sustained microglial activity after FLASH exposure. Meanwhile, CONV-induced genes were mainly involved in iron related homeostasis and membrane/lipid biosynthesis, as well as promotion of proliferation and migration (Fig. 4F). The top enriched pathways uniquely induced after FLASH show enhanced phagocytic and proteolytic activity (*Megf10, Mmp1C*) as well as tissue remodeling (*Adam1S, CdSSl2, Spon2, Col13a1*), showing a microglial phenotype that is prone to ensure tissue clearance. Additionally, the top pathways showing the biggest difference in enrichment score between FLASH and CONV mainly revealed pathways upregulated by CONV and involved in immunomodulatory signals (promoting T cell infiltration *IP10/CXCL10, Il5*), ceramide, UV-repair and eye development (Fig. 4H).

### FLASH modulates the neuronal transcriptome in the hippocampus

Importantly, the radiation-induced signature in neurons was shown to be region-specific. In the DG, FLASH altered the neuronal transcriptome 24h post-irradiation, while the transcriptomic impact of CONV was minimal (Fig. 5A). The 28 genes upregulated by FLASH (Supp. Table 3) reflected a coordinated stress-adaptive response with genes involved in DNA repair (*Ercc1, Bub1, Ube2d2b, Fbxo28*), neuroprotection by detoxification of lipid peroxidation products and oxidative stress mitigation (*Gsta4, Adh1, Pgrmc1*), mitochondrial/proteostasis control (*Mrs2, Gpbar1*), glial/immune activation (*Spib, Gpbar1*), and ECM remodeling (*Eln, Egfem1*). Gene Set Enrichment Analysis (GSEA) showed FLASH-specific upregulation of ATPase regulator activity and chloride/anion and ligand-gated ion channel pathways (Fig. 5B), suggesting an inhibitory, homeostatic signature, limiting excitotoxic damage.

**Figure 5.**
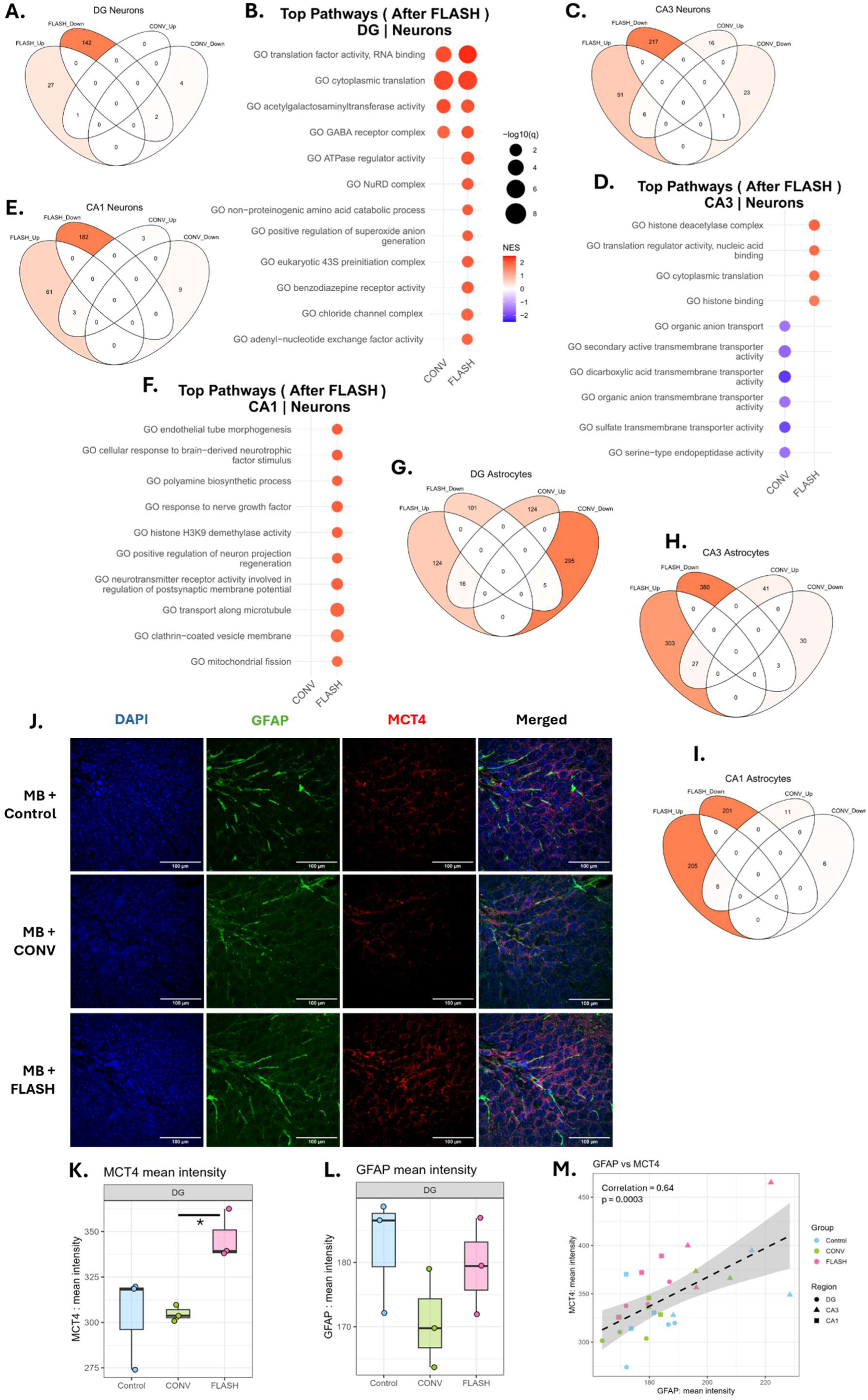
FLASH induces a distinct neuronal and astrocytic transcriptomic imprint in the hippocampus. **A/C/E/G-I-**Venn diagrams of the significantly differentially up- and down regulated genes in the neurons of the DG (A), CA3 (C), and the CA1 (E), and in the astrocytes of the CA1 (G), CA3 (H), and the DG (I) after either CONV or FLASH-RT. **B/D/F-**Gene set enrichment analysis (GSEA) of the neurons in the DG (B), CA3 (D), and the CA1 (F). Pathways are ranked according to the normalized enrichment score (NES). The top pathways enriched after FLASH are displayed. Color represents the NES, where a negative NES means a downregulated pathway. The significance is displayed as the size of the dots. **J-**Representative confocal images of the immunofluorescence co-staining of GFAP (green) and MCT4 (red) in the DG. N=3 mice per group, the same brains as for the DSP were used. Imaged at 100x, scale bars are 100 µm. Images are a projection of the z-stack, using the max intensity, acquired in ImageJ. **K-**Quantification of the MCT4 mean fluorescence intensity per mouse in the DG across the treatment groups (n=3 mice/group; box: median and IQR; points: individual mice). Statistical analysis was done with one-way ANOVA and Tukey’s post hoc test: *p< 0.05. **L-**Quantification of the GFAP mean fluorescence intensity per mouse in the DG across the treatment groups (n=3 mice/group; box: median and IQR; points: individual mice). **M-**Relationship between GFAP and MCT4 mean intensity across all mice and hippocampal subregions (DG, circles; CA3, triangles; CA1, squares), colored by treatment group. Pearson correlation gives r=0.64, p=0.0003.

FLASH upregulated 97 genes in the CA3 (Fig. 5C, Supp. Table 3), amongst which genes important for activity-dependent and synaptic plasticity (*Arc, Camk2g, Adcy4, Calb1, Chrna4, Kcnip2, Adarb1, Pth2r, Pnoc*). Similar to the DG, upregulated genes are involved in mitochondrial metabolism (*Dnm1l, Ckmt2, Plin5*), axonal transport and vesicle trafficking (*Vps2Cb, Gak, Entr1, Bicd1, Bloc1s1, Golga5*), cytoskeletal remodeling (*Dock3, Arpin, Insc, Speg, Fras1, Adam20*), and transcriptional reprogramming (*Emx1, Nkx1-2, Hoxb1, Klf3, Klf15, ZbtbS, Dpy30, Eid3, CnotS*). GSEA revealed FLASH-driven upregulation of chromatin/histone regulation and translation, whereas only CONV suppressed transmembrane transporter activity, suggesting an active repair/plasticity response under FLASH versus a functional impairment under CONV (Fig. 5D).

Lastly, in the CA1, the transcriptional response of the neurons was altered more after FLASH, like in the other hippocampal subregions (Fig. 5E), and 64 genes were upregulated by FLASH (Supp. Table 3). Contrary to the plasticity genes found in the CA3, this signature is more enriched in genes related to stress regulation (*Atn1, Patz1, Ebf3, Bclaf3, Cdca7l, Ctdspl, Foxn1, Bsx*). Interestingly, the upregulation of Esr1 suggests an estrogen-sensitive modulation of plasticity. Other upregulated genes are involved in transcriptional control (*Aplf, Etaa1, Dzip3*), proteostasis (*Ubqln1, Fbxo1C, Fbxo30, Arrdc4*), metabolic remodeling and lipid regulation (*Akr1cC, FabpC, Etnppl, Apool, Mrpl3*), anti-inflammatory genes (*Entpd1/CD3S, Cnr2*), and ion channels (*Asic4, SlcCa5, Slc30a2, SlcSb2*). GSEA showed FLASH-specific enrichment of neurotrophic factor signaling, neuron projection/regeneration, and postsynaptic receptor/vesicle trafficking pathways, consistent with the active pro-survival and plasticity engagement (Fig. 5F).

### FLASH and CONV induce a distinct transcriptomic signature in the astrocytes of the hippocampus

Astrocytes in the different subregions of the hippocampus are functionally heterogeneous, and as observed for the neurons, the radiation-induced transcriptome in astrocytes was found to be dose-rate dependent and region-specific. Interestingly, unlike the neurons, the astrocytes showed a balanced up/downregulation ratio. In the DG, both modalities altered the astrocytic transcriptome largely 24h post-irradiation (Fig. 5G, Supp. Table 3), with FLASH upregulating many astrocytic functions including cell cycle, chromatin protection, and DNA surveillance genes (*Gadd45gip1, Hexim1, Atmin, Bid, Ezh2, Baz1a, Chrac1*), and neuron-glia communication factors (*Il33, Entpd3, Fgf2, Ffar2, Lcn2*).

In the CA3, a major transcriptional impact was caused only by FLASH (Fig. 5H, Supp. Table 3), with 330 upregulated genes that can be clustered into several coherent functional programs: metabolic support (*CoxCa1, Cox8a, NdufsC, Atp5a1, Sod2, Pygb, Arg2*), synapse-glia communication (*Gria1, Grik4, Gabarap, Glrb, Gpr158, Chrm4, Cnr2, Bsn, Vamp2, Sv2b, Sh3gl2*), regulation of inflammatory signaling (*Ccl2, Alox5, Fcgr1, Cd300lb*), and chromatin remodeling (*Hdac2, Ash2l, Rnf2, Elk4, Fosl1, Hey1*).

The CA1 also showed a major transcriptional signature, but only after FLASH (Fig. 5I, Supp. Table 3). FLASH-upregulated genes involved in ECM remodeling (Col1a1, Mxra7, Ctnna1, Itgb5, Gja5, Cldn22), steroid/lipid metabolism (Hsd3b7, Hsd3b3, Abca9, Esrra, Vdr), transcriptional control (Kdm5b, Hoxa1, Foxo6, Hey2, Med21), and synaptic modulation (Gabra1, Kcnk5, Kcng4, Trpc3, Adcy1, Gnas) were identified. Additionally, the upregulated genes suggest a reactive astrocytic phenotype (Tlr4, Nlrc5, Ifng, Ifngr2, Csf2rb, Cxcr3).

Transcriptomic data suggests a neuron-supporting signature induced in astrocytes by FLASH. To support this hypothesis the expression of the astrocytic monocarboxylate transporter MCT4, a key transporter of lactate from astrocytes to neurons, was assessed by GFAP immunofluorescence in the DG, CA3, and CA1 (Fig. 5J, Supp. Fig. 5 A/D). Findings corroborated this hypothesis, as significantly higher MCT4 levels were found 24h after FLASH-irradiation in the DG (Fig. 5K), with a similar trend in the CA3 and the CA1 (Supp. Fig. 5C/F). GFAP expression did not show a significant reduction 24h post-CONV in the DG (Fig. 5L), with a similar trend found in the CA3 (Supp. Fig. 5B) and no differences in the CA1 (Supp. Fig. 5E). Across all mice and hippocampal subregions, GFAP and MCT4 showed a positive correlation (r=0.64, p=0.0003) (Fig. 5M).

## Discussion

This study showed that hypo-fractionated FLASH radiotherapy delivered in 3 fractions of 10 Gy (BED = 60 Gy, α/β=10, standard-of-care), achieved complete and durable control of medulloblastoma in an immuno-compromised mouse model of both sexes, equivalent to CONV dose-rate irradiation. In addition, stable NOR performance was measured in all irradiated animals and was equivalent to age-matched tumor-free non-irradiated animals. However, molecular analysis performed in female mice revealed reduced long-term neuroinflammation in the FLASH cohort, further supported by the results of the early DSP profiling performed across multiple regions-of-interest. Transcriptomic analyses performed in the cerebellum, where the MB cells were engrafted, and in tumor-free hippocampal subregions showed distinct dose-rate-dependent signatures.

FLASH induced substantially more transcriptional changes than CONV across all regions-of-interest, with a different pattern depending on the region. In the cerebellum, a highly divergent transcriptional program was revealed after FLASH versus CONV in the microglia, with minimal overlap between the two modalities. FLASH induced a broader transcriptional shift with 876 upregulated genes supporting sustained metabolic activation and ionic remodeling. This enhanced phagocytic and proteolytic activity is indicative of a microglial phenotype oriented towards debris clearance and tissue remodeling. In contrast, CONV upregulated pathways linked to iron homeostasis, membrane/lipid biosynthesis, proliferation and migration. While our study was performed in adaptive immunocompromised animals, the myeloid contribution remained intact, and our findings are consistent with two recent reports showing a differential role for the microglial cells after FLASH vs CONV ^17, 22^. In diffuse midline glioma, Padilla et al. showed that electron irradiation in both FLASH and CONV was able to activate an IFN1 response, while early activation of microglial cells occurred uniquely after FLASH, consistent with our results. Ni et al. used proton FLASH irradiation in a transgenic medulloblastoma (Math1-Cre/smoM2+/-) model, showing that it abrogates lipid oxidase expression and oxidized low-density lipid generation to reduce PPARγ activity ^17^. Meanwhile, CONV induced reactive oxygen species-dependent PPARγ, resulting in differential and immunosuppressive polarization of tumor-infiltrating macrophages. Interestingly, in our study, the pathways exhibiting the greatest differential enrichment between CONV and FLASH were predominantly CONV-driven. They included proliferative and immunomodulatory signaling cascades dependent upon IP10- and IL5-pathways after CONV, while FLASH promoted a metabolically active, phagocytosis-prone state potentially supportive of efficient tissue clearance able to shape a distinct inflammatory milieu. Despite these distinct acute signatures, both modalities ultimately resulted in equivalent and durable tumor response, suggesting that the high dose regimen selected was ablative and drove the therapeutic outcome.

The hippocampal signatures obtained from three ROIs located in functionally linked and sequential areas including the DG, which is essential for memory encoding; the presynaptic CA3, essential for pattern separation; and the post-synaptic CA1, essential for pattern completion, were distinct ^23, 24^. FLASH was able to modify the transcriptional program of astrocytes at 24h post-RT, with genes aligning closely with the pattern of each region to support preserved neuronal function ^25^. In the DG, astrocytes exhibited signatures related to metabolic support, lipid handling, and extracellular matrix modulation, suggesting enhanced neuron–glia metabolic coupling to sustain plasticity as proposed previously ^26, 27^. In addition, the up-regulation of *Gadd45gip1, Hexim1, Atmin, Bid, Ezh2, Baz1a, Chrac1* suggest that FLASH-exposed astrocytes are activating cell cycle control, chromatin protection and DNA surveillance programs but not overt pro-apoptotic collapse prone to maintain tissue stability. The upregulation of *Il33, Entpd3, Fgf2, Ffar2, Lcn2* suggest active neuron–glia communication, fine-tuned immune response and trophic support signaling. In the CA3, astrocytes showed upregulated genes involved in oxidative phosphorylation, cytoskeletal remodeling, membrane trafficking, and immune-related signaling, indicative of a metabolically active, synapse-engaged, and dynamically responsive glial state. Enhanced metabolic fitness, including mitochondrial respiratory chain components (*CoxCa1, Cox8a, NdufsC, Atp5a1*), antioxidant defense via Sod2, and energy metabolism genes such as *Pygb* and *Arg2* were found. This metabolic reinforcement was paralleled by enrichment of synapse–glia communication pathways, including glutamatergic signaling (*Gria1, Grik4*), GABAergic components (*Gabarap, Glrb*), GPCR-mediated neuromodulation (*Gpr158, Chrm4, Cnr2*), and synaptic scaffolding/vesicle dynamics genes such as *Bsn, Vamp2, Sv2b, and Sh3gl2*, suggesting preserved excitatory–inhibitory balance to promote efficient neurotransmission, as shown in another study ^28^. Interestingly, no evidence of a broad pro-inflammatory cascade was found, indeed the upregulation of *Ccl2, Alox5*, *Fcgr1*, *Cd300lb* suggested a controlled immune reaction. Finally, the upregulation of chromatin modifiers including *Hdac2, Ash2l, Rnf2* and activity-linked transcription factors such as *Elk4, Fosl1,* and *Hey1* support sustained transcriptional reprogramming that has been shown to support neuronal plasticity and cognitive function ^29^. Strikingly, CA1 astrocytes displayed a robust extracellular matrix, interferon/immune, lipid–steroid metabolic, and chromatin remodeling profile, consistent with structural stabilization and long-term modulation of the synaptic milieu at the principal node of hippocampal output. The increased MCT4 expression found in astrocytes of the DG after FLASH irradiation reinforces the hypothesis that astrocytes support neuron function. Furthermore, such observations align with the oxidative phosphorylation and energy metabolism signatures identified after FLASH and identify possible astrocyte-to-neuron metabolic coupling through the lactate shuttle, a topic that warrants further investigation.

Collectively, data is consistent with astrocytic function in the DG, that helps to maintain firing required for pattern separation ^30^, in the CA3, that helps protect neurons against ischemia by suppressing intracellular Ca^2+^ overload ^31^ and in the CA1, that regulates synaptic plasticity and information integration ^32^. A recent study also provides evidence for a robust astrocyte to neuron conversion upon inhibition of PTBP1 in a model of Alzeimer’s disease ^33^ worthy for future FLASH experimentation.

Concomitantly, in the neurons of the DG, FLASH-induced upregulation was enriched with genes involved in cell cycle–associated chromatin dynamics, DNA repair, mitochondrial function, and metabolic adaptation. This is consistent with heightened structural plasticity and activity-dependent remodeling in granule cells. Mitochondrial and metabolic function seemed sustained through upregulation of *Dnm1l,* known to control mitochondrial fission ^34^ and through *Ckmt2,* a mitochondrial creatine kinase required to maintain energy homeostasis ^35^. *Plin5,* known to decrease the exposure of mitochondria to fatty acid by storing them in lipid droplets ^36^, was also found to be upregulated in the CA3. In the CA3, FLASH modulated neuronal transcripts prominently reflect synaptic organization, vesicle trafficking, and excitatory transmission, while CONV induced only modest changes. The neuronal signature largely involved the activity-dependent plasticity axis, including *Arc* ^37^*, Camk2g* ^38^*, Adcy4* ^39^*, and Calb1* ^40^ supporting preserved synaptic plasticity. This aligns with the pattern-separation properties of the CA3 and is consistent with the synaptic vesicle recycling and long-term potentiation (LTP) signature found recently in tumor-free adult mice exposed to FLASH ^41^. The concurrent upregulation of mitochondrial and metabolic genes such as *Dnm1l, Ckmt2,* and *Plin5* ^34, 36^ indicates reinforced bioenergetic capacity, while enrichment of genes involved in axonal transport (*Bicd1)* ^42^, vesicle trafficking in the brain (*Vps2Cb, Gak*) ^43, 44^, cytoskeletal remodeling supporting axonal growth (*Dock3)* ^45^, and dendritic spine morphology *(Arpin*) ^46^ suggest structural adaptability. Upregulation of genes involved in transcriptional reprogramming in neurons (*Klf3, Klf15)* ^47, 48^ and *Dpy30* known to regulate epigenetic landscape in neural progenitors ^49^ suggest a sustained activity-dependent gene regulation. The signatures of the neurons in the CA1 shifted toward synaptic output regulation, intracellular signaling cascades, and transcriptional control pathways, possibly associated with memory consolidation-related plasticity. This is consistent with previous spatial transcriptomic profiling, where memory consolidation-related transcriptional programs in CA1 neurons were predicting and functionally modulating long-term memory formation ^50^. Several anti-inflammatory genes were also found, including *Entpd1***/***CD3S,* a phosphatase known to dampen extracellular ATP signaling and therefore limiting inflammation ^51^ and *Cnr2,* a receptor known to modulate neuronal excitability with potential anti-inflammatory action ^52^, as well as ion channels and transporters (*Asic4, SlcCa5, Slc30a2, SlcSb2)* involved in the maintenance of ionic balance ^53^.

Together, these findings support a model in which FLASH does not induce a uniform activation program but rather drives circuit-tailored astrocyte-neuron crosstalk (Figure 6), as described by Vanrobaeys upon learning tasks in young mice ^50^. Regulatory changes induced by FLASH in the DG would favor plasticity initiation, enhance recurrent network support and metabolic resilience in the CA3, and promote structural consolidation and immune-modulated synaptic refinement in the CA1. This coordinated, region-specific reprogramming suggests that FLASH engages distributed astrocyte-neuron partnerships to reshape hippocampal circuit function at multiple hierarchical levels leading to long-term cognitive preservation. Aside from the molecular imprints specific to changes in dose rate, equivalent therapeutic outcomes suggest further that hypo-fractionated radiotherapy regimens may represent a clinically attractive strategy to reduce the logistical and psychological burden associated with multiple treatment sessions, particularly in pediatric and frail patient populations.

**Figure 6.**
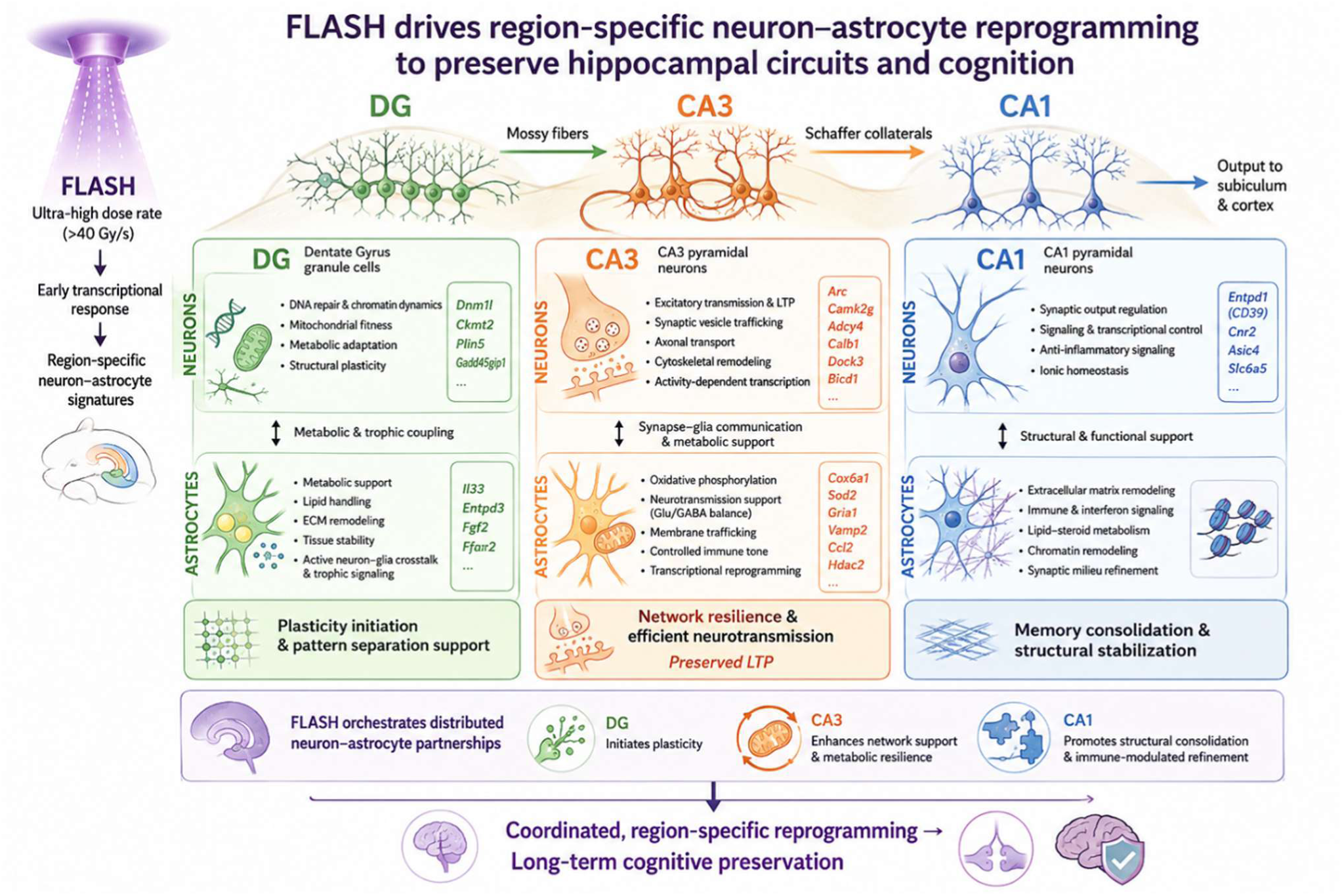
FLASH drives region-specific neuron-astrocyte reprogramming to preserve hippocampal circuits and cognition. Schematic overview of the region- and cell type-specific transcriptomic effects of FLASH irradiation on several hippocampal subregions. Summary figure generated using ChatGPT.

As with most studies, conclusions need to be evaluated alongside certain caveats and limitations. First, the intermediate-energy electron FLASH irradiator limits penetration and is not conformal, precluding targeted treatment of deep-seated brain tumors but enabling the use of high dose-rate which remains today the gold standard for FLASH preclinical studies (> 10^6^ Gy/s). Investigating whether lower dose rates (100 Gy/s) provided by proton and photon FLASH irradiators would induce similar molecular responses remains to be validated. Although one single cell line/tumor was used and sex representation was not fully balanced, our study did show similar tumor response and survival outcomes in male and female without significant differences in behavioral performance between CONV-RT and FLASH-RT conversely to our previous publication ^10, 14^. In these previous works, immunocompetent juvenile mice were used while here the immunocompromised status of the animals is certainly the major confounding factor impacting the neurocognitive testing alongside with orthotopic tumor implantation.

In conclusion, the present data supports the concept that FLASH irradiation induces qualitatively distinct biological responses that may promote medulloblastoma clearance and mitigate radiation-induced neurotoxicity. Optimization of dose, fractionation, radiation modality, and conformal delivery will be the critical next steps to fully harness the therapeutic window of FLASH radiotherapy and translate its potential cognitive-sparing benefits to children with brain tumors.

## Acknowledgments

We acknowledge the statistical expertise of Dr. Craig Stark from UCI for his invaluable support. We want to thank Prof. Martin Baumgartner – University Children’s Hospital Zürich authorized by Dr. John Silber C Michael Bobola from Seattle Children’s Hospital and Regional Medical Center for their UW228 human MB cells donation that made this work possible. We thank the histology core facility at UNIGE, specifically Marie Napoli, for cutting the samples. Finally, we would like to thank the core facilities and staff at Lausanne CHUV, as well as their veterinary staff, for their valuable support.

## Author Contributions

Conceptualization and experimental design: PBZ, CLL and MCV. Methodology and experimental development: JFP, MK, AA, LVK, BP, AJ, JO, JJ, VG, JR and PBZ. Data analysis: MK, JFP, AA, LK, BP, AJ, JO, CL, MCV and PBZ. Writing, review and editing: All. Project administration and funding acquisition: PBZ, CLL and MCV.

