## Supplementary material for "Differential and regionally distinct molecular responses distinguish FLASH and conventional RT in a medulloblastoma model while preserving therapeutic outcomes": supp data

#### Supplementary Material and method

##### GeoMx Digital Spatial Profiling at an acute time point

Brain samples were fixed in 4% PFA and embedded in paraffin. 5 µm sagittal sections were placed on a slide with one CTRL, one CONV, one FLASH per slide. Spatial transcriptomics was performed according to the standard nanoString GeoMx Digital Spatial Profiler (DSP) protocol[1], using the Whole Transcriptome Atlas (WTA) panel. Specific markers were used to identify different cell types: neurons (NeuN, 1:100, Millipore ABN78, with a secondary goat anti-rabbit Alexa 594, 1:1000, Abcam ab150080), astrocytes (GFAP Alexa 532, 1:100, Novus NBP2-3318), and microglia (CD68 Alexa 647, 1:200, Abcam ab305214), as well as a nuclear marker (Syto83, Invitrogen 511364).

###### *Bioinformatics*

Quality control, pre-processing, and further analysis of the data was performed in R (v4.4.0) as per the recommendations from nanoString[2], using custom scripts and publicly available packages, apart from the normalization of the data, which was done using the Trimmed Mean of M-values (TMM) method. Out of the 110 initial segments, 4 were removed during the quality control because of a low number of nuclei (<50) or aligned reads (<75%), 17 were removed during the detection rate filtering as fewer than 5% of the genes had an expression above the limit of quantification. In summary, quantification from 89 robust segments was possible for downstream analysis. Specificity of the cell type annotations was evaluated by a marker gene expression heatmap, showing concordance with known cell type-associated markers (Supp. Fig. 3A). To assess the sensitivity and coverage of the sequencing, gene detection rates were examined across all cell types and regions, showing acceptable detection rates in all cell types and regions, with higher detection rates in the neurons and the cerebellum (Supp. Fig. 3B). To evaluate possible batch-driven errors, dimensionality reduction using a uniform manifold approximation and projection (UMAP) was performed and did not reveal any clustering (Supp. Fig. 3C and 3D). Together, these analyses confirmed data suitability for downstream analyses.

#### Immunofluorescence at an acute timepoint

Immunofluorescence was used to evaluate the level of GFAP and MCT4 in the hippocampus 24h after radiotherapy (n=3 per group). Brain sections were prepared as described above. The following primary antibodies were used: rabbit anti-MCT4 (1:100, AMT-014, Alomone) and sheep anti-GFAP (1:200, AF2594, Bio-Tech), in combination with the secondary antibody Alexa Fluor 568 donkey anti-rabbit (1:500, A10042, Life Technologies) and Alexa Fluor 488 donkey anti-sheep (1:500, A11015, Life Technologies). Slides were mounted with Prolong anti-fade with DAPI to stain the nuclei. Images were acquired on a confocal microscope (Nipkow) at a 100x using 5 z-slices with 1 µm separation. Analysis was performed using the QuPath software for acquiring the intensities and ImageJ for acquiring the z-projection. The mean intensity was determined from the three middle z-slices using a full image annotation and was afterwards averaged over the three slices.

#### Evaluation of neuroinflammation at endpoint

Six mice of each group sampled six months after radiotherapy by intracardiac perfusion with PBS and brain was processed to analyze neuroinflammation.

***Total RNA*** was isolated from longitudinal sections of PFA-fixed and paraffin-embedded brain tissues using the MagMAX™ FFPE DNA/RNA Ultra Kit (Applied Biosystems, USA) (n=4 per group). The concentration of RNA isolated was measured using a NanoDrop spectrophotometer (Thermo Fisher, USA). 0.5 µg of RNA from each sample was used to synthesize cDNA using the SuperScript™ VILO™ cDNA Synthesis Kit (Invitrogen, USA). Quantitative real-time polymerase chain reaction (RT-qPCR) was performed using TaqMan™ Fast Advanced Master Mix and different TaqMan® Gene Expression Assays (Applied Biosystems, USA). All genes were assessed in duplicate and our assays contained Template Controls as negative controls containing all the components of the PCR master mix but without cDNA, which was substituted with an equivalent volume of RNase free water. We examined changes in the expression of the inflammatory markers *Il-1b, Tnf-a, Il-6, Nlrp3, Il-4* and *Gsdmd*. Gene expression levels were normalized to endogenous controls *Hprt* and *Gapdh* and calculated using the 2−ΔΔCt method. Assays ID used are listed in Supp Table 1.

***Immunofluorescence*** was used to evaluate microglial activation and astrogliosis in the cerebellum and hippocampus (n=4 per group). Primary antibodies were as follows: rat anti-Iba1 (1:200, ab283346, Abcam, England), rabbit anti-CD68 (1:200, ab283654,

Abcam, England), rabbit anti-GFAP (1:200, PA5-16291, Invitrogen, USA). Secondary antibodies were Alexa Fluor 594 goat anti-rabbit IgG (1:500, 111-585-144, Jackson ImmunoResearch, USA) or Alexa Fluor 488 goat anti-rat IgG (1:500, 112-545-16, Jackson ImmunoResearch, USA). Nuclei were visualized by incubating with DAPI. Images were acquired on a Nikon A1R+ inverted confocal microscope using NIS Elements C software and ImageJ analysis (% area positive).

Normality of the datasets was assessed using the Shapiro–Wilk test and confirmed by visual inspection of Q–Q plots prior to parametric statistical analyses. Since data met normality assumptions, results were analyzed using a one-way ANOVA and Tukey's post hoc test for multiple comparisons between treatments.

#### Supplementary Table.

**Supplementary Table 1. Irradiation parameters**

| Beam Parameters | CONV | FLASH |
|---|---|---|
| | 3 x 10 Gy | 3 x 10 Gy |
| Graphite applicator type and size (mm) | Hemi-circular Ø17 | Hemi-circular Ø17 |
| Source-to-surface distance (mm) | 800 | 209 |
| Pulse repetition frequency (Hz) | 10 | 100 |
| Pulse width | 1.0 µs | 1.8 µs |
| Number of pulses | 1170-1180 | 1 |
| Treatment time (s) | 117 | $1.8 \cdot 10^{-6}$ |
| Mean dose rate (Gy/s) | 0.1 | $5.6 \cdot 10^{6}$ |
| Instantaneous dose-rate (Gy/s) | $8.5 \cdot 10^{3}$ | $5.6 \cdot 10^{6}$ |

**Supplementary Table 2.** Taqman probe details (Applied Biosystems, USA) used for RT-qPCR.

| Target gene | NCBI Reference Sequence | Taqman Assay ID | Product size |
|---|---|---|---|
| *Il-1b* | NM_008361.3 | Mm00434228_m1 | 90 |
| *Tnf-a* | NM_001278601.1 | Mm00443258_m1 | 81 |
| *Il-6* | NM_031168.1 | Mm00446190_m1 | 78 |
| *Nlrp3* | NM_145827.3 | Mm00840904_m1 | 84 |
| *Il-4* | NM_021283.2 | Mm00445259_m1 | 79 |
| *Gsdmd* | NM_026960.4 | Mm00509958_m1 | 94 |
| *Hprt* | NM_013556.2 | Mm03024075_m1 | 131 |
| *Gapdh* | NM_001289726.1 | Mm99999915_g1 | 107 |

**Supplementary Table 3.** List of up- and down-regulated genes per cell type and subregion.

#### Supplementary Figures

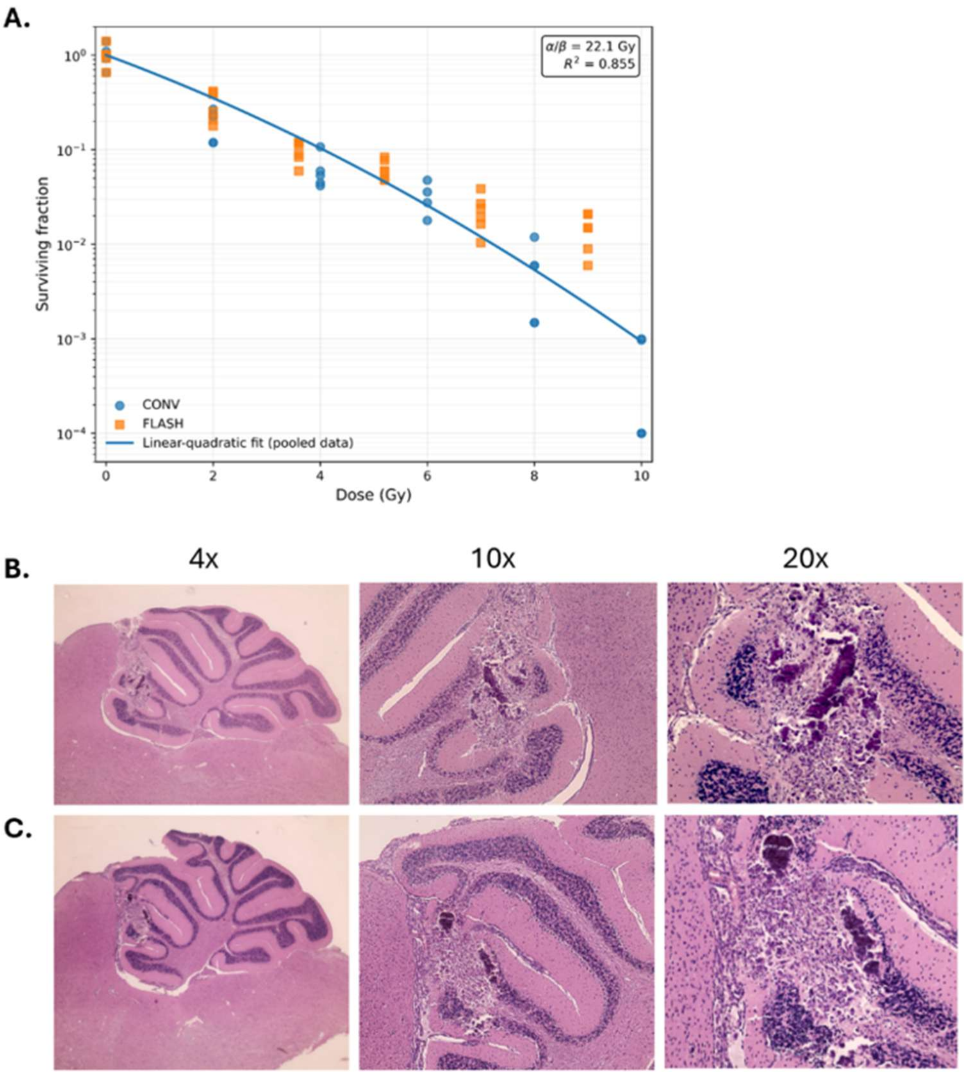

**Supplementary Figure 1. A-** Estimation of the α/β ratio of the UW228-2 medulloblastoma cell line using the linear-quadratic model. Individual clonogenic surviving fractions obtained after conventional and FLASH irradiation were pooled to estimate the intrinsic radiosensitivity of the cell line. Data points represent individual biological replicates after exclusion of predefined outliers. Surviving fractions were log-transformed and fitted according to the linear-quadratic model, with the intercept constrained to zero (SF=1 at 0 Gy). The fitted parameters were $\alpha=0.480\ Gy^{-1}$, $\beta=0.0217\ Gy^{-2}$, corresponding to an estimated α/β ratio of 22.1 Gy. **B-C Histological confirmation of tumor presence at the time of radiotherapy.** Representative H&E-stained cerebellar sections from two female UW228-2 tumor-bearing mice collected 24 h after irradiation, corresponding to approximately 2 weeks after orthotopic tumor implantation. Representative sections from two independent animals shown at 4×, 10×, and 20× magnification. Histological examination demonstrates tumor tissue at the implantation site despite the absence of radiographically detectable lesions by contrast-enhanced micro-CT at the time of treatment.

**A.**

| Groups | Cohorts | | Total mice and number of Batches | | Valid for Behavioral Test | |
|---|---|---|---|---|---|---|
| | Females | Males | Females | Males | Females | Males |
| Tumor Free 0Gy | Batch0( n=5)<br>Batch5 (n=10) | | n=15<br>2 batches | | n=10 | |
| MB 0Gy | Batch 2 (n=8)<br>Batch 3 (n=8) | Batch 1(n=4)<br>Batch 7(n=5)<br>Batch 9 (n=10) | n=16<br>2 batches | n=19<br>2 batches | n=8 | n=8 |
| MB+CONV 3x10Gy | Batch 1(n=4)<br>Batch 2 (n=4)<br>Batch 3 (n=4)<br>Batch 7 (n=4) | Batch 1(n=4)<br>Batch 8 (n=16) | n=16<br>4 batches | n=20<br>2 batches | 4 months n=14<br>6 months n=12 | 4 months n=19<br>6 months n=19 |
| MB+FLASH 3x10Gy | Batch 1(n=4)<br>Batch 2 (n=4)<br>Batch 3 (n=4)<br>Batch 7 (n=4) | Batch 8 (n=16) | n=16<br>4 batches | n=16<br>1 batches | 4 months n=16<br>6 months n=10 | 4 months n=15<br>6 months n=15 |

**B.**

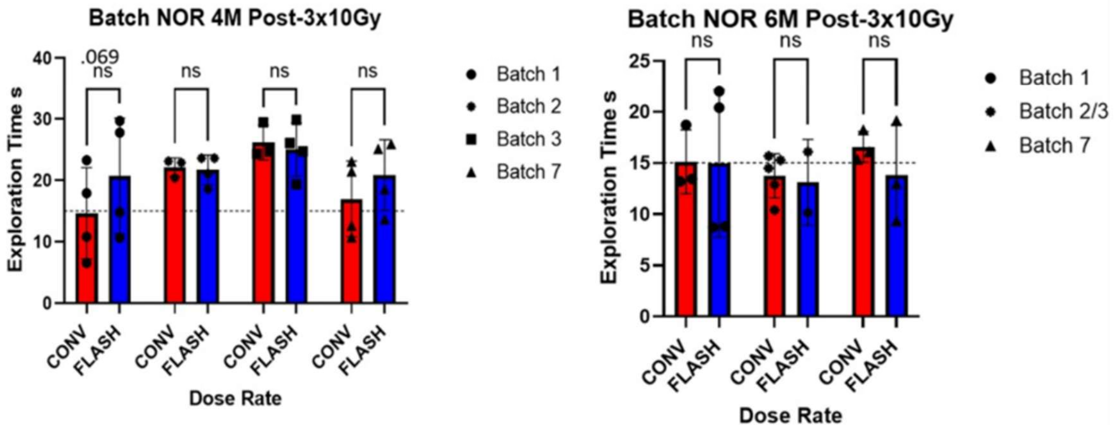

**Supplementary Figure 2. Experimental cohort composition and batch-specific analysis of novel object recognition. (A)** Experimental cohort composition showing the total number of animals assigned to each experimental group, their distribution by sex, and the number of independent experimental batches. Cohort sizes represent the animals included in each experimental cohort and may therefore exceed the number contributing to individual behavioral analyses because of data availability and statistical exclusion of outliers. **(B)** Novel-object exploration during the first 30 s of cumulative object exploration shown separately for the independent experimental batches of the female cohort. Individual values are shown and batch-specific analyses are presented to illustrate the consistency of the behavioral observations across independent experimental cohorts.

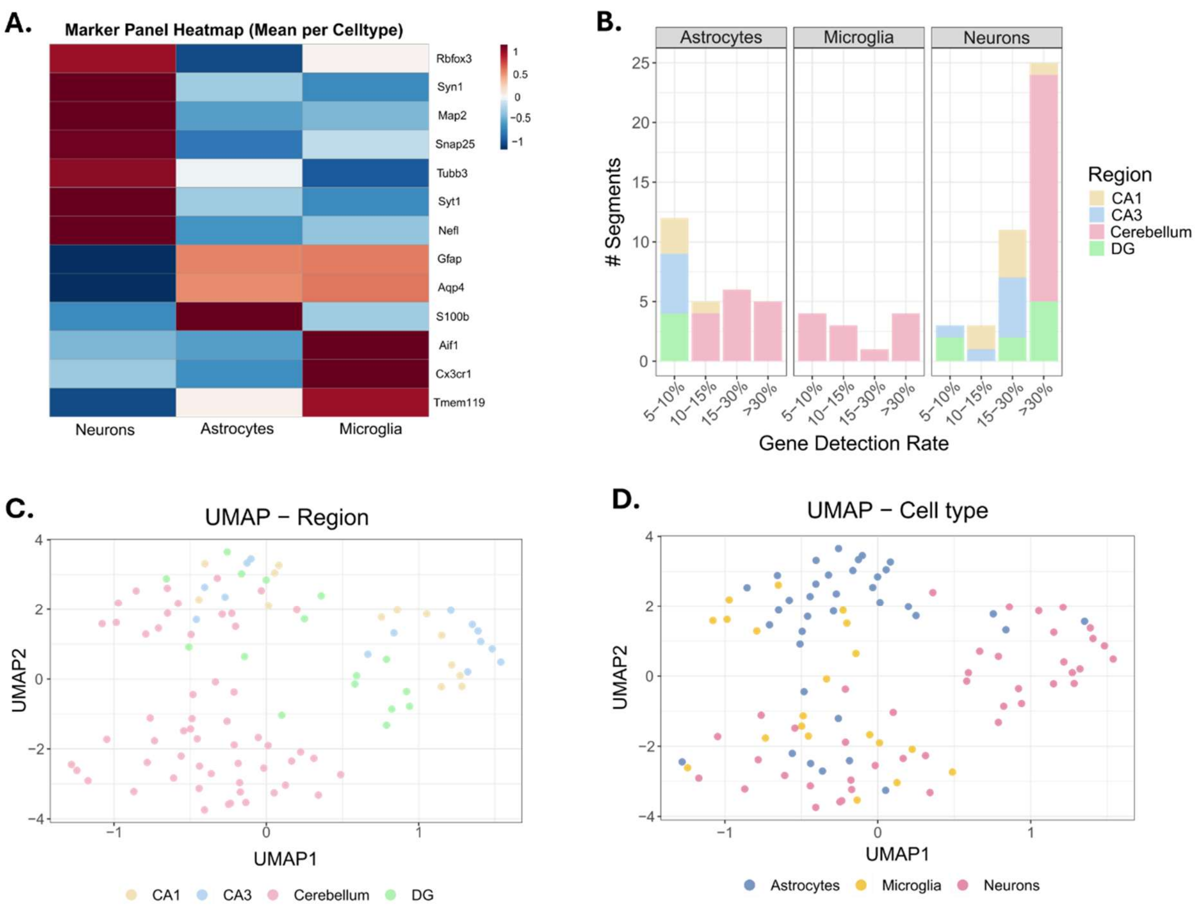
A.
Marker Panel Heatmap (Mean per Celltype)
Rbfox3
Syn1
Map2
Snap25
Tubb3
Syt1
Nefl
Gfap
Aqp4
S100b
Aif1
Cx3cr1
Tmem119
Neurons
Astrocytes
Microglia
B.
Astrocytes
Microglia
Neurons
### Segments
Gene Detection Rate
5–10%
10–15%
15–30%
>30%
Region
CA1
CA3
Cerebellum
DG
C.
UMAP − Region
UMAP1
UMAP2
CA1
CA3
Cerebellum
DG
D.
UMAP − Cell type
UMAP1
UMAP2
Astrocytes
Microglia
Neurons

**Supplementary Figure 3. A-** A heatmap showing the expression of a marker panel per cell type. Normalized expression values per cell type were summed over all regions and normalized per row to obtain expression values between -1 and 1. **B-** Stacked barplot showing the gene detection rate per region and cell type. **C-** UMAP dimensionality reduction plot showing all segments (n = 89), colored by region. **D-** UMAP dimensionality reduction plot showing all segments (n = 89), colored by cell type.

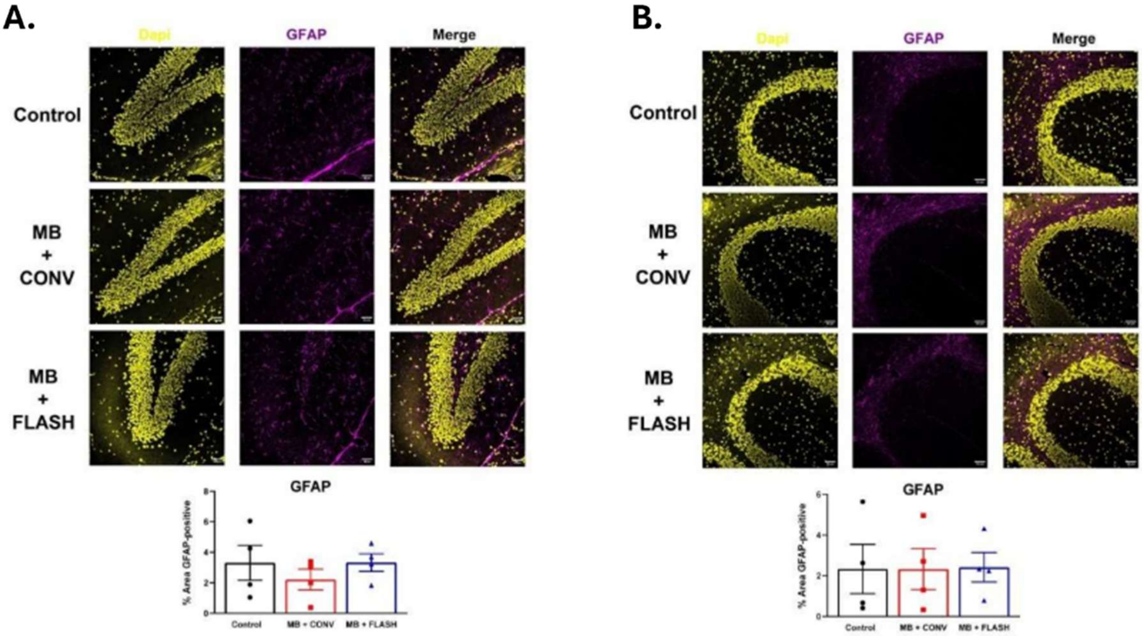

A.
Dapi
GFAP
Merge
Control
MB + CONV
MB + FLASH
GFAP
% Area GFAP-positive
Control
MB + CONV
MB + FLASH
B.
Dapi
GFAP
Merge
Control
MB + CONV
MB + FLASH
GFAP
% Area GFAP-positive
Control
MB + CONV
MB + FLASH

**Supplementary Figure 4. A-, B-** No significant changes were observed in the hippocampus (A) and cerebellum (B) when we analyzed GFAP as an astrogliosis marker. Data are expressed as mean ± SEM (n = 3-4 per group). The analysis was done with one-way ANOVA and Tukey's post hoc test. *$p < 0.05$, **$p < 0.01$, ***$p < 0.001$.

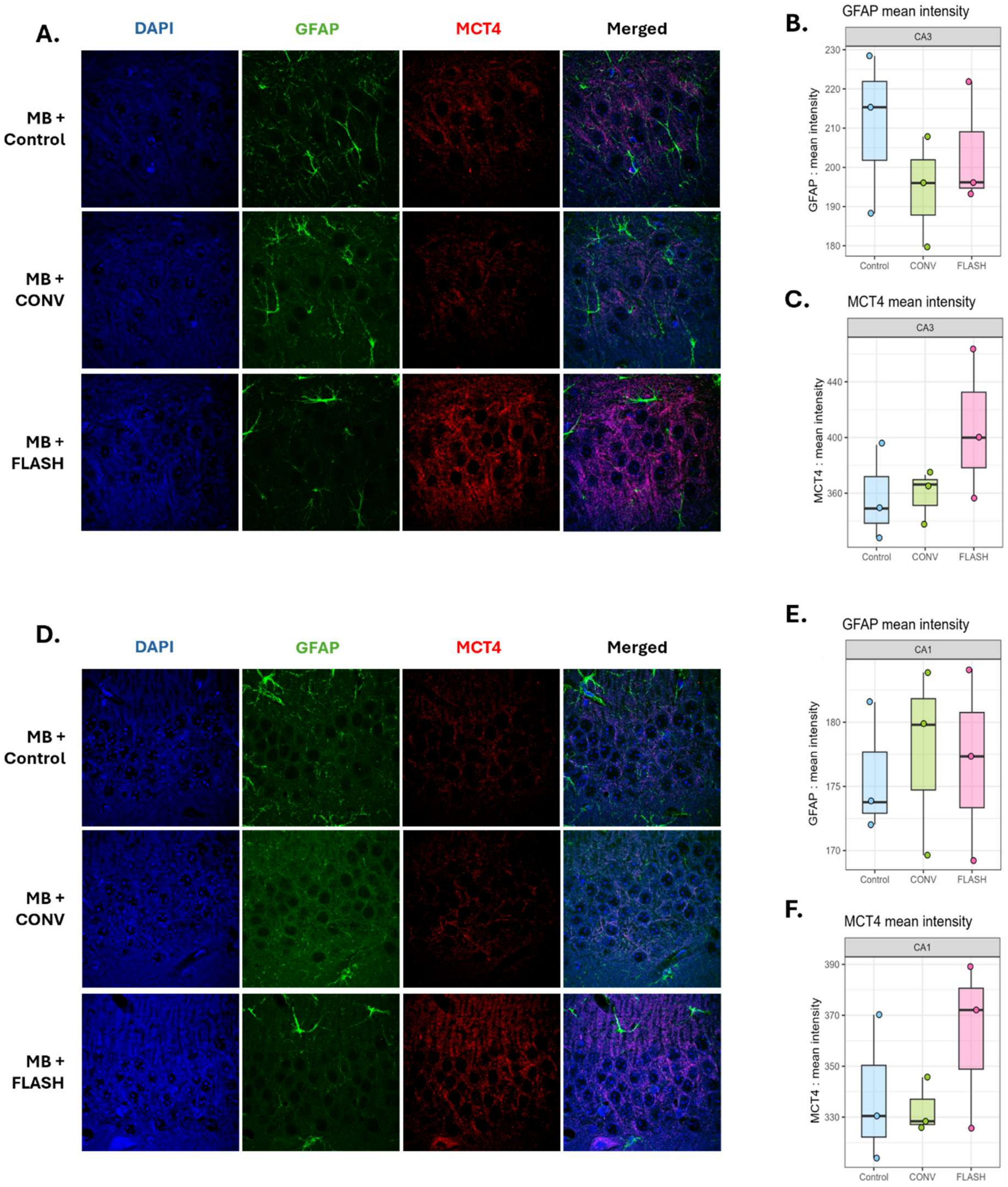
A.
DAPI
GFAP
MCT4
Merged
MB +
Control
MB +
CONV
MB +
FLASH
B.
GFAP mean intensity
CA3
GFAP : mean intensity
230
220
210
200
190
180
Control
CONV
FLASH
C.
MCT4 mean intensity
CA3
MCT4 : mean intensity
440
400
360
Control
CONV
FLASH
D.
DAPI
GFAP
MCT4
Merged
MB +
Control
MB +
CONV
MB +
FLASH
E.
GFAP mean intensity
CA1
GFAP : mean intensity
180
175
170
Control
CONV
FLASH
F.
MCT4 mean intensity
CA1
MCT4 : mean intensity
390
370
350
330
Control
CONV
FLASH

**Supplementary Figure 5. A-D-** Representative confocal images of the immunofluorescence co-staining of GFAP (green) and MCT4 (red) in the CA3 (A) and the CA1 (D). n=3 mice per group, the same brains as for the DSP were used. Imaged at 100x, scale bars are 100 μm. Images are a projection of the z-stack, using the max intensity, acquired in ImageJ. **B-E-** Quantification of the GFAP mean fluorescence intensity per mouse in the CA3 (B) and the CA1 (E) across the treatment groups (n=3 mice/group; box: median and IQR; points: individual mice). Statistical analysis was done with one-way ANOVA and Tukey's post hoc test. **C/F-** Quantification of the MCT4 mean fluorescence intensity per mouse in the CA3 (C) and the CA1 (F) across the treatment groups (n=3 mice/group; box: median and IQR; points: individual mice). Statistical analysis was done with one-way ANOVA and Tukey's post hoc test.
